# Discovery and Visualization of Age-dependent Patterns in the Diurnal Transcriptome of Drosophila

**DOI:** 10.1101/2022.04.06.487011

**Authors:** Benjamin Sebastian, Rosalyn M. Fey, Patrick Morar, Brittany Lasher, Eileen S. Chow, Jadwiga M. Geibultowicz, David A. Hendrix

## Abstract

Many critical life processes are regulated by input from 24-hour external light/dark cycles, such as metabolism, cellular homeostasis, and detoxification. The circadian clock, which helps coordinate the response to these diurnal light/dark cycles, remains rhythmic across lifespan; however, rhythmic transcript expression is altered during normal aging. To better understand how aging impacts diurnal expression, we present an improved Fourier-based method for detecting and visualizing rhythmicity that is based on the relative power of the 24-hour period compared to other periods (RP24). We apply RP24 to transcript-level expression profiles from the heads of young (5-day) and old (55-day) *Drosophila melanogaster*, and reveal novel age-dependent rhythmicity changes that may be masked at the gene level. We show that core clock transcripts phase advance during aging, while most rhythmic transcripts phase delay. Transcripts rhythmic only in young flies tend to peak before lights on, while transcripts only rhythmic in old peak after lights on. We show that several pathways including glutathione metabolism, gain or lose coordinated rhythmic expression with age providing insight into possible mechanisms of age-onset neurodegeneration. Remarkably, we find that many pathways show very robust coordinated rhythms across lifespan, highlighting their putative roles in promoting neural health. We investigate statistically enriched transcription factor binding site motifs that may be involved in these rhythmicity changes.

## Introduction

Numerous cellular processes, including energy metabolism, homeostasis and detoxification, are coordinated according to 24-hour cycles due to input from external light/dark cycles and regulation by the internal circadian system. Circadian control of many diurnally-expressed genes imposes temporal coordination on signaling and enzymatic pathways leading to optimal organismal functions. Maintenance of circadian control prevents metabolic dysregulation [1] and appears critical for healthy neuronal aging and longevity; both flies and mice with disrupted circadian clocks are prone to accelerating aging and are more susceptible to neurodegeneration and oxidative stress [2-5]. Most core clock genes are rhythmically expressed in young animals and these rhythms may be altered with age in a tissue dependent manner; however, molecular clock oscillations continue in old organisms, indicating that clocks are functional [6-8]. Aging organisms may show diurnal gene expression in response to light which is perceived as stress, especially in the blue part of the spectrum [9]. Blue light exposure induces oxidative stress, which results in gene expression changes including the upregulation of stress-response genes [10]. Studies of blue-light-exposure in aging flies reveal the induction of age-specific stress response genes, increased neurodegeneration, and reduced lifespan [11].

While the clock remains functional across lifespan, recent studies have revealed age-dependent changes in the expression patterns of clock-controlled genes, which act downstream from the clock to regulate cellular processes. Analysis of human postmortem brain samples revealed substantial differences in age-associated rhythmic gene expression, including genes that gained rhythmicity with age [12]. Similarly, increased rhythmicity was reported in studies comparing age-related changes in gene expression in several mouse tissues [6, 7]. We sequenced and analyzed around-the-clock RNA from 5-day (young) and 55-day (old) *white Drosophila melanogaster* heads and found that aging fly tissues express a new set of genes in a circadian or diurnal manner, which we named late life cyclers (LLCs) [8]. While these reports focused on surprising age-induced gene oscillations, pathways that lose or maintain rhythmicity with age did not receive adequate attention.

To address these complex age-related changes in transcript expression, we developed a systems-level characterization of the diurnal transcriptome to identify rhythmic pathways that change with age and those that remain intact across lifespan. Current methods for detecting rhythmically expressed transcripts rely on computational techniques that identify expression profiles resembling a sinusoid or a similar pattern, such as Fourier analysis [13], goodness of fit to a sine wave [12], non-parametric statistical tests [14, 15], and harmonic regression [16]. Here, we created and utilized a simple yet robust signal-to-noise ratio, relative power of the 24-hour period (RP24), that builds upon previous Fourier-based approaches [13] to identify transcripts with the most robust rhythms and to characterize their rhythmicity changes during aging. We identified large-scale changes in transcriptomic rhythmicity and phase in aged flies, investigated functional enrichment and explored candidate regulators of these groups.

## Results

### Detecting oscillatory transcriptomic patterns

Fourier-based approaches to detect diurnal expression patterns typically examine the discrete Fourier power spectrum, which breaks down time series information into the frequency domain and its inverse, the Fourier periods. For transcript expression at *N* discretely sampled time points over a duration *T*, the power spectrum *P*(*T*_*k*_) quantifies the strength of the periods *T*_*k*_ = *T*/*k* for *k* = 1, …, *N* − 1. A common approach for identifying rhythmic expression profiles is to examine the *F*_24_: the power of the 24-hour Fourier period relative to the average value for random permutations of the transcript expression time series [13, 17].

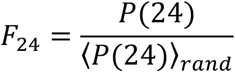

Because biological data is inherently noisy, expression profiles with a strong 24-hour component often have deviations from a smooth oscillation that result in strong components for non-24-hour Fourier periods. Therefore, we devised a score that accounts for this by computing the relative power of the 24-hour period compared to all other periods (RP24). The expression profile for any transcript *E*(*t*) can be decomposed into a 24-hour component and noise terms, such that:

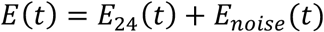

We assume the noise term *E*_*noise*_(*t*) is only composed of non-24-hour periodicity because any 24-hour periodic oscillations in the noise would be absorbed into *E*_24_(*t*), the 24-hour component. We then consider the ratio of the total power of *E*_24_(t) and the total power of *E*_*noise*_(*t*), which is the signal-to-noise ratio:

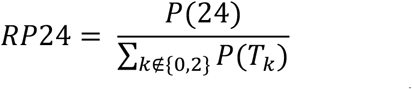

In this expression, *P*(*T*_*k*_) is the discrete Fourier power spectrum of *E*(*t*), and *T*_*k*_ = 48/*k* are the periods of the Fourier decomposition. The only non-zero term of the power spectrum of *E*_24_(*t*) is the 24-hour component, and the *E*_*noise*_(*t*) only has non-24-hour components of the power spectrum. The RP24 is a score that quantifies the 24-hour signal in a transcript expression profile relative to noise; therefore, changes in the RP24 value for a group of transcripts should indicate a change in the fidelity of 24-hour signals in the associated expression profiles.

We compared the RP24 to the *F*_24_ for each transcript in young flies (Figure 1A). While these scores are correlated, the upward curvature of the scatter plot demonstrates the ability of the RP24 to separate highly rhythmic transcripts compared to the *F*_24_. For example, the two most rhythmic expression profiles observed in young flies are the completely uncharacterized transcripts *CG16798-RA* and *CG44195-RA* (Figure 1B). The power spectrum for each transcript exists only as a 24-hour component (Figure 1C), but while the *F*_24_ value for these two transcripts is similar to many other transcripts, the RP24 better highlights them as highly-rhythmic genes. In contrast, the transcripts *pain-RA* and *Nup54-RA* were selected as examples that are not rhythmic according to RP24, but have large *F*_24_ values. (Figure 1D and 1E). The transcript *pain-RA* has an *F*_24_ of 3.5, but moderate 16-and 12-hour components result in a much lower RP24 of 1.7. The transcript *Nup54-RA* has an *F*_24_ greater than 2, while the strong 9.6-hour component results in an RP24 of 0.54. The presence of non-24-hour components in the expression profiles of these two transcripts cause a less-precise rhythm that deviates more dramatically from a smooth sinusoidal curve, reflected in the lower RP24 score compared to the *F*_24_.

**Figure 1:**
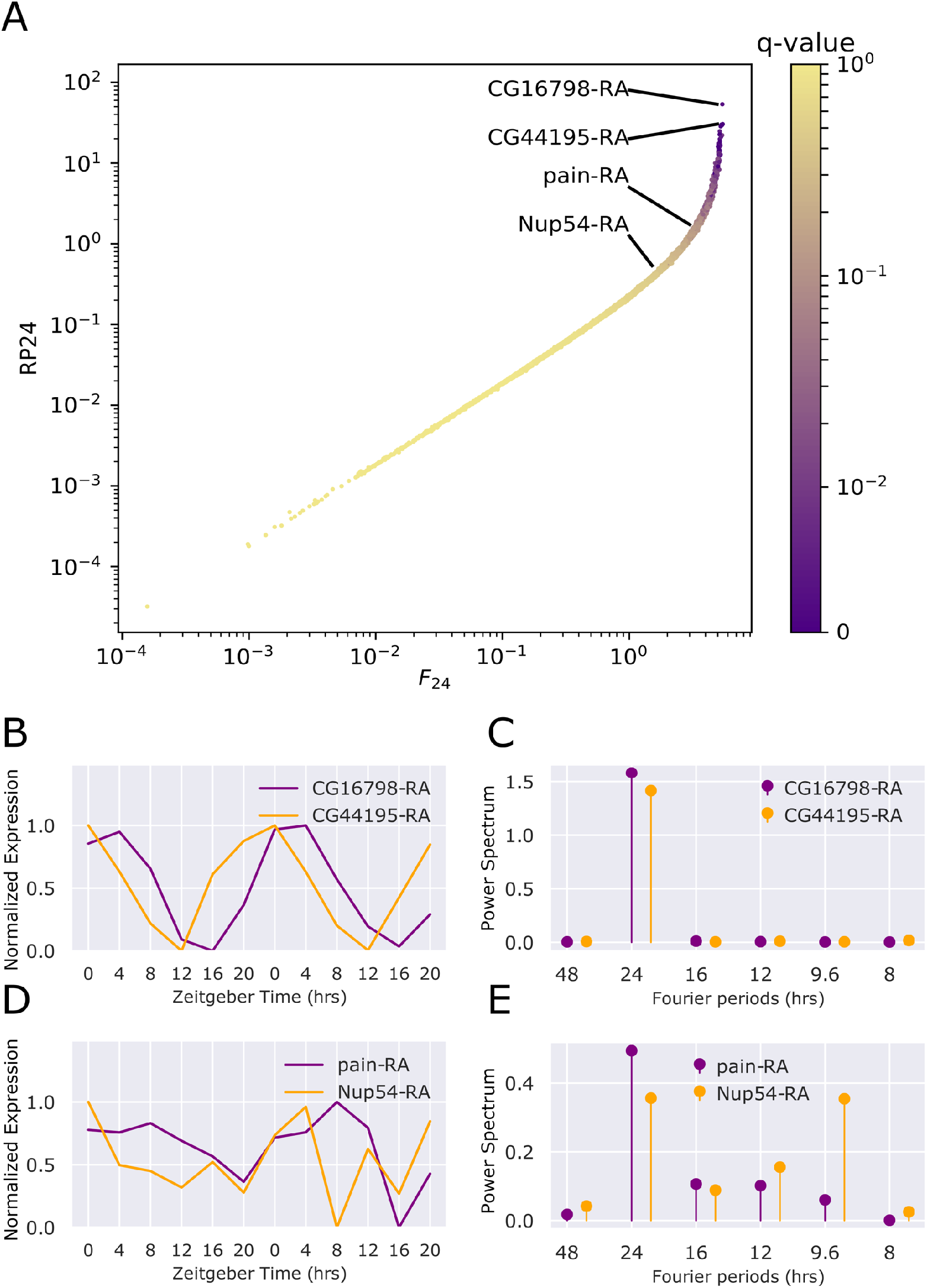
Comparison of RP24 score to *F*_24_ score. **A**. Scatterplot comparing the RP24 score to *F*_24_, defined as the fold change of the power of the 24-hour component of the expression profile over random permutations. **B**. The expression profiles of the two most rhythmic transcripts, CG16798-RA and CG44195-RA in our young (5-day) flies. **C**. The Fourier power spectrum of the two most rhythmic transcripts in young flies. **D**. The expression profiles in young flies of the two least rhythmic transcripts based on the RP24 q-value, *pain-RA* and *Nup54-RA*, that have a *F*_24_ score greater than 3. **E**. The Fourier power spectrum of *pain-RA* and *Nup54-RA* in young flies.

### Widespread transcript-specific alterations of rhythmic expression patterns associated with aging

Using the RP24, we evaluated changes in rhythmicity for each transcript in young and old flies. We compared the distribution of RP24 values over all transcripts between young and old (Figure 2A) and found a statistically significant (p-value = 3.4e-9) increase in net rhythmicity in old flies using a Kolmogorov-Smirnov test.

**Figure 2:**
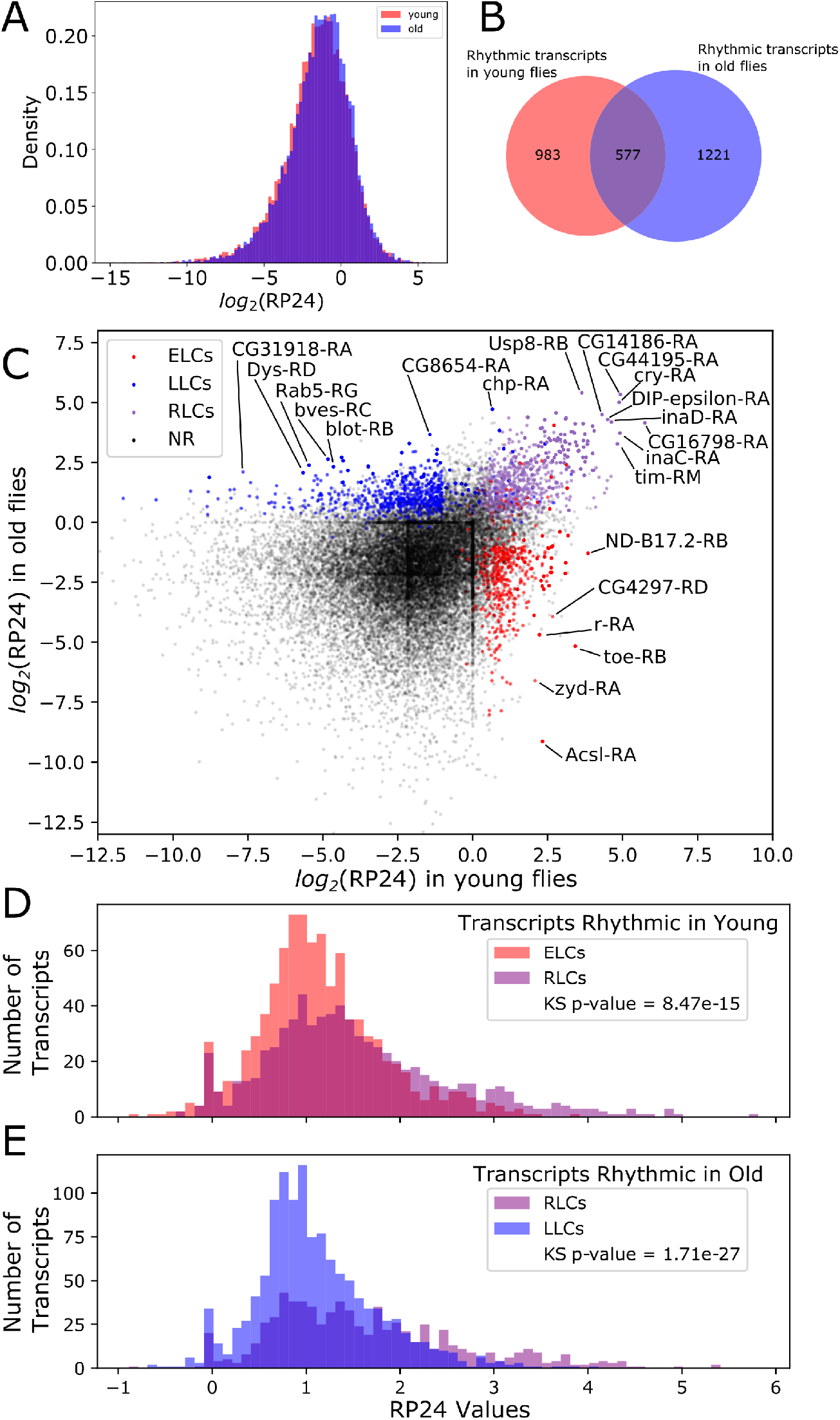
Rhythmicity changes with age. **A**. The total amount of rhythmicity, as defined by the distribution of RP24 over all transcripts, changes slightly toward greater rhythmicity after aging.**B**. Euler diagram shows that the specific transcripts with statistically significant RP24 values is substantially different between young and old flies. **C**. A scatterplot where each transcript is represented by a dot. The x-axis value is the log-transformed RP24 value in young flies, and the y-axis position is the log-transformed RP24 value in old flies. Red dots correspond to transcripts that are significantly rhythmic (FDR≤0.05) in young flies and not in old, blue dots correspond to transcripts that are significantly rhythmic in old flies and not young, and purple dots correspond to transcripts that are rhythmic in both young and old flies. **D**. Histogram shows the RP24 distribution for ELC compared to RLC transcripts in young flies. **E**. Histogram shows the distribution of RLC compared to LLC transcripts in old flies.

The difference in the RP24 distributions between young and old fly transcriptomes is explained by numerous individual transcripts that gain or lose rhythmicity with age. To identify statistical changes in rhythmicity at the transcript level, we computed a p-value for each transcript by comparing its RP24 value to the distribution of RP24 values generated by randomly shuffling the time points of its expression profile. We performed a Benjamini-Hochberg multiple test correction on the resulting p-values to define q-values for each transcript.

To observe age-associated trends, we divided the rhythmicity continuum into discrete rhythmicity states defined by q-value range. Transcripts were considered rhythmic if they had a statistically significant RP24 value (q-value ≤ 0.05) and arrhythmic if they had a q-value ≥ 0.075. Indeterminant transcripts (0.05 < q-value < 0.075) were filtered out of our analysis. We also filtered transcripts with periodic spikes of expression because these could not be validated experimentally (see Methods). We defined “detectable” rhythmic transcripts as those having at least 1.5-fold change between maximum and minimum expression (max/min fold change) and a median expression level of at least 1 FPKM [8, 18, 19]. Using these criteria, we identified 1560 detectable rhythmic transcripts in young flies and 1798 detectable rhythmic transcripts in old flies. The overlap between young and old (577 transcripts) are transcripts rhythmic in both young and old flies (Figure 2B).

We used these parameters to define broad groups of age-dependent transcript expression changes, similar to our previous work [8]. We identified 742 early life cyclers (ELCs), which are transcripts that show statistically significant rhythmicity in young flies but are arrhythmic in old flies. We observed 1024 late life cyclers (LLCs), which are rhythmic in old flies but arrhythmic in young flies. To account for borderline cases of transcripts that are rhythmic in both ages, we require robust life cyclers (RLCs) to be detectable in one age, but allow a lower max/min fold change of 1.4 in the other, which results in 628 RLCs. Among the RLCs we found the core clock transcripts, *Clk-RA, tim-RB, tim-RM, tim-RO, per-RA, Pdp1-RJ, Pdp1-RD, Pdp1-RP, vri-RA* and *vri-RE*, indicating that the circadian clock remains rhythmic with age. Unexpectedly, we also detected rhythmicity in the cycle gene (*cyc-RA*), which has been considered the only clock gene with no discernable cycling [20, 21]. In addition to clock transcripts, RLCs contained transcripts derived from 105 genes that were previously identified as clock-controlled in heads of young flies [17], supporting the notion that the circadian clock remains functional with age. Complete lists of all transcripts in each group are provided in Supporting Table S1. Figure 2C compares RP24 in young (x-axis) and old (y-axis) flies. Each transcript is represented by a dot with a color corresponding to the rhythmicity group, and with “not rhythmic” (NR) indicating transcripts not in the ELC, RLC or LLC groups.

We next used RP24 distributions to compare the level of rhythmicity for the ELCs, RLCs and LLCs. We found that in young flies, the RLCs have on average greater rhythmicity than ELCs (Figure 2D). In old flies, RLCs have on average greater rhythmicity than LLCs (Figure 2E). In both cases, the RLC RP24 histograms have a longer tail, and the difference of the two distributions is statistically significant by a Kolmogorov-Smirnov-test.

In addition to the RP24 score to detect rhythmic transcripts with a 24-hour period, we defined a similar score to identify transcripts oscillating with other periods *T*. We found that ELCs in old had an increase in 12-hour periods (Supporting Figure S1). Similarly, we found an increase in 9.6-hour periods for LLCs in young flies.

### Phase changes after aging

Transcript expression profiles with the most significant RP24 values will have well-defined, precise rhythms for which phase can be reliably calculated. We generated simulated transcript expression profiles to test accuracy of phase calculations, and found that the phase calculations are accurate, and most accurate for larger RP24 values (Supporting Figure S2A-B). We calculated the phase for each transcript for both young and old expression profiles (see Methods) and observed age-related phase changes in rhythmic expression. We used circular histograms plotted over a 24-hour clock to visualize the phase distributions for young and old expression profiles for the RLCs (Figure 3A). The general trend is toward a phase delay after aging, with peak expression time shifting from before lights-on (ZT0) to after lights-on. We also compared the phases of ELCs in young flies to the phases of LLCs in old flies (Figure 3B). We observed a substantial difference in the two distributions that was consistent with the trend for the RLCs: a phase shift from before to after lights-on. The median phase for the ELCs was ZT23.31 (0.69 hours before lights on), while the median phase for the LLCs in old was ZT2.4 (2.4 hours after lights-on).

**Figure 3.**
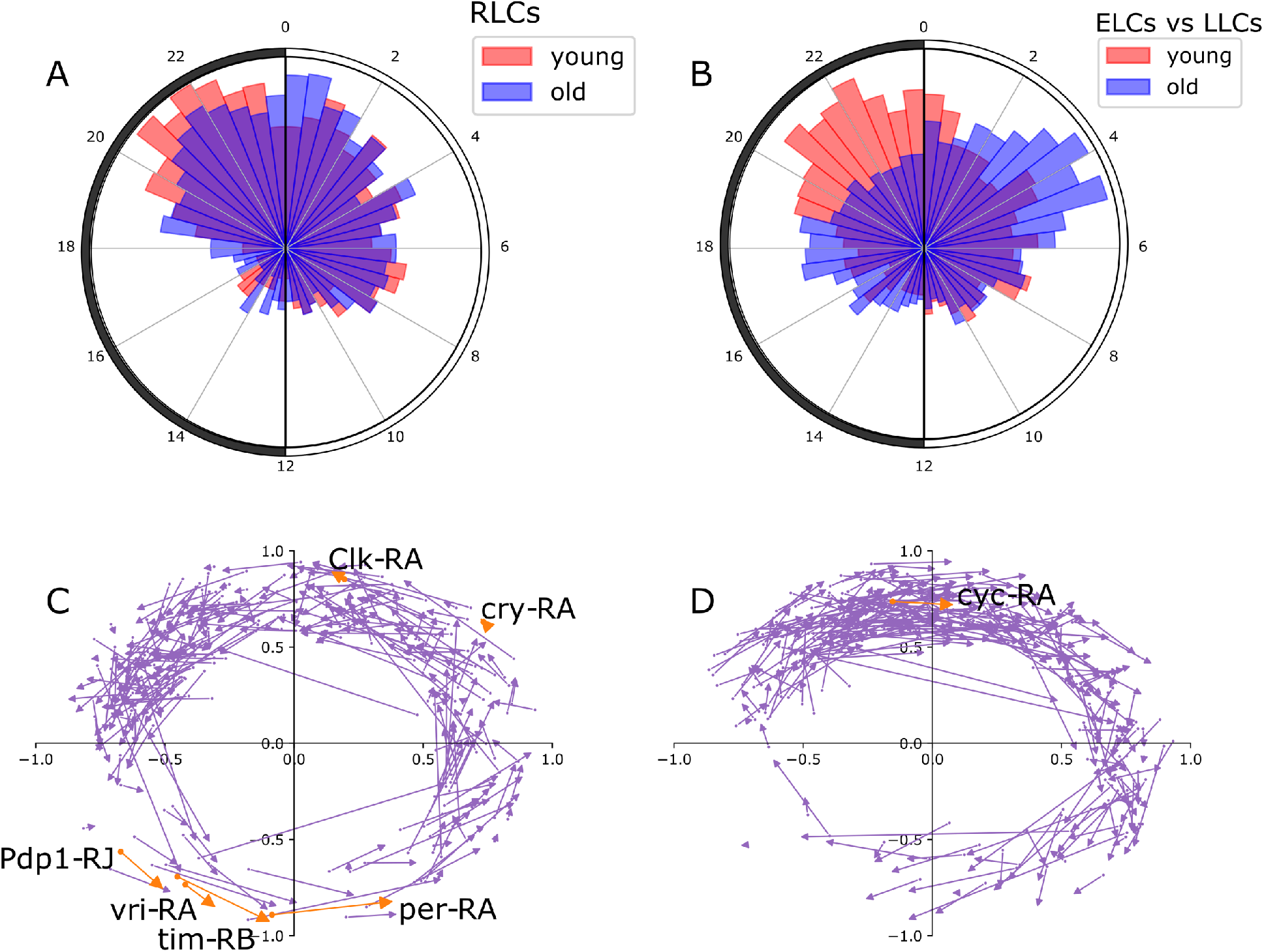
Phase changes with age. **A**. Circular histogram depicts the phase of RLC transcripts in young compared to the distribution of the phase of the same transcripts in old. The light-dark cycles is represented on the histogram as a 24-hour clock, with light-on (ZT0) at the top and lights-off (ZT12) at the bottom. Phases are binned into increments of 30 minutes. **B**. Circular histogram similar to panel C compares the phase distribution of ELC transcripts in young to LLC transcripts in old. **C**. Dot-and-arrow scatterplot shows phase advance for RLC transcripts. Distance from the origin is to the dot is RP24 in young, scaled between 0 and 1 with a logistic function, and the phase is shown as the angle from the positive y-axis (ZT0) to the phase of that gene on the same 24-hour clock as panels C and D. Each transcript is represented moving from a phase/rhythmicity in young (scatter point) to a phase/rhythmicity in old (tip of arrow). Transcripts belonging to the core clock mechanism are shown in orange and labeled. **D**. Dot-and-arrow scatterplot shows phase delay for RLC transcripts.

We created “dot-and-arrow” scatterplots to visualize age-associated changes in rhythmicity that shows changes in phase as well as RP24. The state of rhythmicity of each transcript is represented by the angle with the positive y-axis (ZT0) in the same 24-hour clock as 3A-B, and the distance from the origin describes the RP24. Figure 3C shows the phase-advanced diurnal expression of each RLC transcript in young represented by a dot; the arrow points to the state of rhythmicity of each transcript in old flies. Figure 3D shows phase-delayed diurnal expression of RLC transcripts. The comparison of Figures 3C and 3D shows that there are more age-dependent phase delays for individual transcripts, as seen by the denser cluster in 3D. Notably, and in contrast to the global trends, the core clock transcripts predominantly exhibit phase advances, with the phase shifting to earlier in the 24-hour cycle (Figure 3C). These differences between subsets of rhythmic transcripts prompted us to further explore alterations in specific subsets during aging.

### Pathway analysis of transcript groups

We assessed whether transcripts in each rhythmicity category were enriched for specific biological pathways using DAVID Functional Annotation Tool [22, 23]. The full list of pathway analysis results is available in Supporting Table S2. Figure 4A shows the enriched pathways identified separately for ELCs, RLCs and LLCs, with color indicating the enrichment score, and the size of each dot corresponding to the number of transcripts in that cluster. The most significant RLC pathways included phototransduction (rhabdomere), mitochondrial translation, locomotor rhythm, choline kinase, and glycolysis. The ELCs showed enrichment of transcripts involved in glutathione metabolism, protein kinase activity, and mitochondrion/transit peptide. The LLC group was enriched for transcripts encoding proteins containing the Pleckstrin homology-like domain (PHD), ATP-binding, and transmembrane domains.

**Figure 4.**
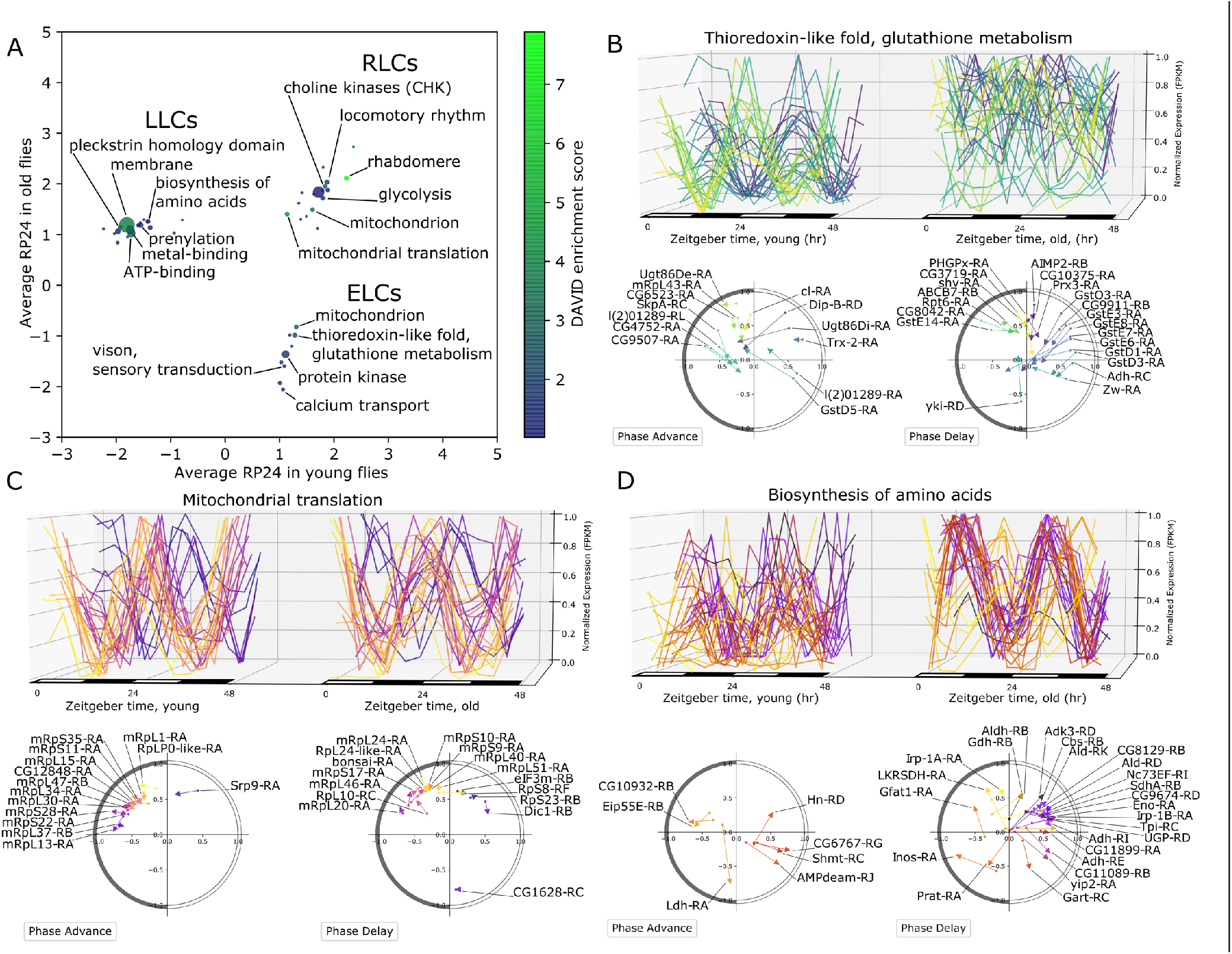
Pathway analysis of rhythmic transcript groups. **A**. Scatterplot representing differentially rhythmic pathways. The x- and y-axes represent the average rhythmicity in young and old, respectively, for DAVID clusters computed from ELC, RLC, and LLC transcripts. The color map represents the DAVID enrichment score, and the size of each dot is proportional to the number of transcripts in that cluster. **B**. The ELC transcripts with thioredoxin-like fold and glutathione metabolism related function show increased expression on average, but with reduced rhythmicity in young compared to old flies. Phase/rhythmicity dot-and-arrow scatterplots (as in Figure 3E-F) are shown for transcripts from this pathway exhibiting phase advance and phase delay. Color map defines a unique color for each transcript based on phase ordering. **C**. The RLC transcripts with mitochondrial translation function show consistent rhythmicity in young and old with little change in phase. **D**. Rhythmicity and phase changes of LLC transcripts with function related to amino acid synthesis.

We further investigated several pathways in each group using our visualization strategies. The ELC pathway that includes glutathione metabolism transcripts shows a clear loss of coordinated expression (Figure 4B). In young, the glutathione-metabolism-associated transcripts can be grouped into two primary phases, but in old the coordinated phases are unrecognizable. The dot-and-arrow scatterplots show that glutathione S-transferases start with a phase between ZT2 and ZT6 in young, but collectively exhibit a phase delay and loss of rhythmicity (Figure 4B). The RLC pathway that includes mitochondrial translation transcripts shows consistent phases between ZT18 and ZT22 in each age, and a very small phase delay with aging (Figure 4C). Lastly, the group of LLC transcripts encoding several enzymes with dehydrogenase activity gained rhythmicity with phases predominately from ZT2 to ZT6 (Figure 4D). This group, which was also annotated with the KEGG pathway term “Biosynthesis of amino acids”, included Alcohol dehydrogenase (*Adh-RI*), Aldehyde dehydrogenase (*Aldh-RI*), Glutamate dehydrogenase (*Gdh-RB*), Succinate dehydrogenase subunit A (*SdhA-RA*), and Lactate dehydrogenase (*Ldh-RA*).

Another group of strikingly rhythmic transcripts is involved in vision, phototransduction, and rhabdomere. We observed that while some transcripts that encode vision-related proteins lose rhythmicity with age (Figure 5A), many others continue to show strong rhythmicity across lifespan (Figure 5C). In addition to this loss of coordination in rhythmic expression, this group of rhythmic transcripts shows varied phase changes with age. Dot-and-arrow scatterplots in Figure 5B and 5D show that approximately equal numbers of transcripts in these groups display phase advance or phase delay during aging. In some cases, different splice variants of the same vision-associated gene can fall into different rhythmicity groups. For example, the gene transient receptor potential like (*trpl*), which encodes a plasma membrane cation channel that is enriched in photoreceptors, has an isoform *trpl-RA* that shows rhythmicity in both ages, while *trpl-RB* is only rhythmic in young (Figure 5B, phase delay). Similarly, the gene neither inactivation nor afterpotential C (*ninaC*), encodes a protein with serine/threonine kinase and myosine activity that is required for photoreceptor function. Transcripts for *ninaC* show divergent age-related rhythmicity, with *ninaC-RA* rhythmic in only young and *ninaC-RD* rhythmic in both young and old. The protein encoded by *ninaC* also forms a complex with *rtp*, which shows a similar phase delay in old while maintaining rhythmicity. This cluster also included sensory transduction transcripts that show different patterns of age-dependent rhythmicity. For example, odorant-binding protein 99a (*Obp99a)* has an LLC transcript, *Obp99a-RA*, while the transcript for the homologous gene *Obp99b, Obp99b-RA* is an ELC. Similarly, *Rh5-RA* has LLC expression, while *Rh3-RA* and *Rh4-RA* are ELCs.

**Figure 5.**
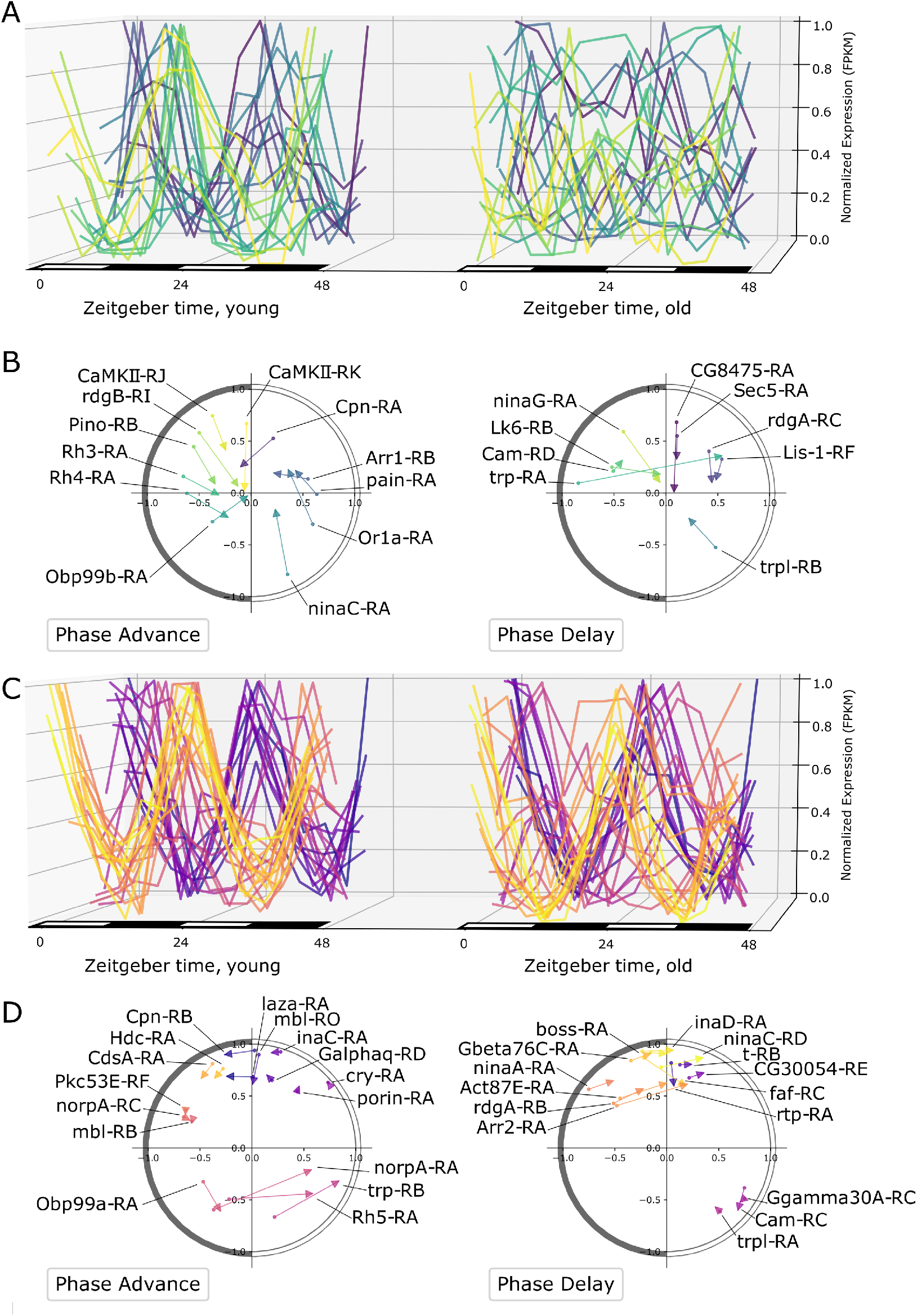
Vision and rhabdomere related transcripts show diverse changes in rhythmicity. **A**. The expression profiles of ELC transcripts related to vision comparing the same transcripts in young and old flies. The color of each curve corresponds to phase ordering in young. **B**. Dot-and-arrow scatterplots show the change in phase and RP24. As with Figure 3E-F, the phase is shown as the angle from the positive y-axis (ZT0) on a 24-hour clock. The color corresponds to the same phase ordering for the same transcripts as panel A. **C**. The expression profiles for RLC transcripts related to vision, comparing young and old. **D**. Dot-and-arrow scatterplots show the change in phase and RP24, similarly as panel B, for the same transcripts in panel C. The color map corresponds to phase ordering in young for both panel C and D.

### Pathway analysis of differentially expressed transcript groups

We performed differential expression analysis to identify transcripts which are significantly up-or down-regulated with age independent of time of day (q-value ≤ 0.05). Of the 3421 differentially expressed transcripts we identified, 647 (18.9 %) belonged to one of the rhythmicity groups defined above. Among the transcripts significantly upregulated in old flies compared to young, we found 101 ELCs, 86 RLCs, and 134 LLCs. We found 88 ELCs, 119 RLCs and 119 LLCs to be significantly downregulated with age. The full lists of up-and down-regulated transcripts for each rhythmicity category are listed in Supporting Table S3.

To understand how up-and down-regulated rhythmic transcripts are involved in aging, we performed pathway analysis on each group of differentially expressed transcripts belonging to ELC, RLC and LLC groups. These results are shown in Figure 6, where the log fold change in transcript expression and the RP24 are averaged over the transcripts in each cluster. A more positive log fold change corresponds to transcripts upregulated in old flies, while a negative log fold change indicates downregulation in old flies.

**Figure 6.**
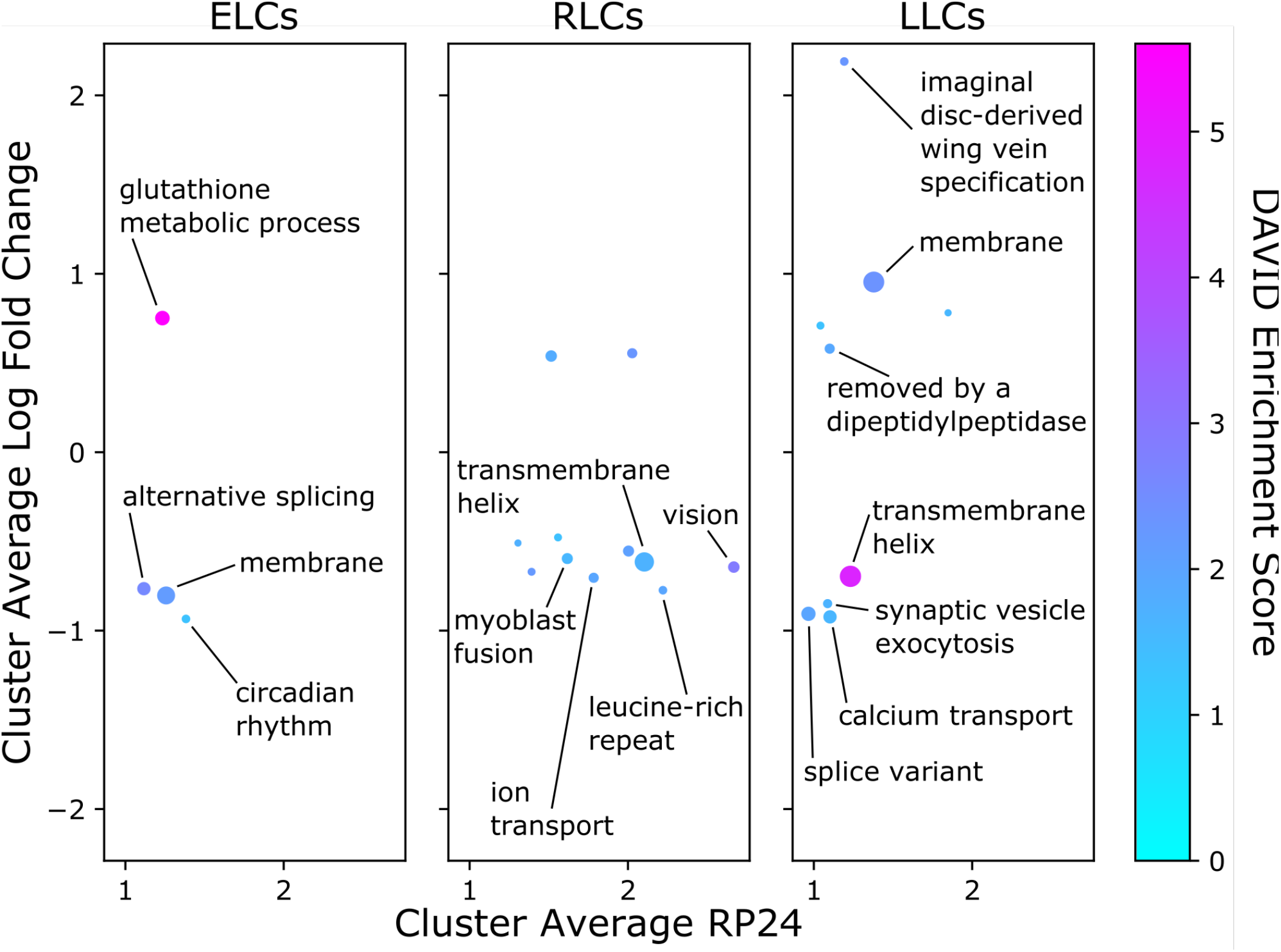
Results of pathway analysis of differentially expressed rhythmic transcript groups. The y-axis of each point corresponds to the log fold change of average expression in old over young for all transcripts in each cluster; the x-axis shows the RP24 averaged over all transcripts in each cluster. Hue denotes DAVID enrichment score, and the size of each point corresponds to the number of transcripts in the cluster. Labeled clusters have at least 5 transcripts, an absolute value log fold change ≥ 0.75, and an enrichment score ≥ 1.3.

The only significant pathway in the upregulated ELC transcripts was glutathione metabolism, including eight transcripts encoding glutathione-S-transferases; however, there are other noteworthy upregulated ELCs. Functional clustering identified twelve ELC transcripts involved in cellular response to DNA damage, including *Pepck-RA, Trxr-1-RB* and *Trx-2-RA*. Consistent with this result, we found among upregulated ELCs three isoforms of Xrp1 (*Xrp1-RG, Xrp1-RC* and *Xrp1-RD*), which is critical for DNA breakage repair [24], and has been shown to be upregulated in response to blue light [10]. The top pathways for upregulated RLCs were flavonoid biosynthetic process and oxidoreductase, the latter of which included the transcript *Eip71CD-RG*, which extends lifespan when overexpressed [25] (Supporting Table S4). We found five enriched pathways for upregulated LLC transcripts, with the most enriched containing transcripts encoding membrane proteins. Upregulated LLCs included *Ldh-RA* and *Hsp26-RA*, both of which are involved in lifespan determination [26, 27].

Downregulated ELC transcripts were enriched in transcripts involved in alternative splicing and transcripts encoding membrane proteins. Also enriched were transcripts playing roles in metabolism, including *Ilp2-RA*, the only isoform of the gene encoding the *Drosophila* Insulin-like peptide 2, knock-out of which has been shown to extend lifespan [28]. Nine significantly enriched pathways for downregulated RLCs included transcripts encoding proteins containing a leucine-rich repeat, and transcripts involved in vision, including *Cpn-RB*, which plays a role in protecting photoreceptor cells from light-induced degeneration [29]. Downregulated LLCs were enriched for splice variant, synaptic vesicle exocytosis, and calcium transport pathways, and transcripts containing a transmembrane helix. The full list of pathway analysis results is available in Supporting Table S4.

While most previous studies of rhythmicity were done at the gene level, our approach shows the utility of analyzing at the transcript level. Indeed, we detected multiple instances where specific transcripts for the same gene showed different age-dependent rhythmicity, which could be masked at the gene-level. For example, *trpl*, an eye-enriched gene involved in photoreceptor response to light, is an RLC at the gene level; however, it has three significantly downregulated isoforms each belonging to a different rhythmicity group (Supporting Figure S3). Another example is the gene *PyK*, encoding pyruvate kinase and involved in glycolysis and glucose metabolism, which has two RLC isoforms, one of which is downregulated with age (*PyK-RA*) and one of which is upregulated with age (*Pyk-RB*) (Supporting Figure S4). We found an even more extreme example in *Gpdh1*, the gene coding for glyercol-3-phosphate dehydrogenase that is involved in triglyceride metabolism. *Gpdh1-RC* is a downregulated LLC and *Gpdh1-RF* is a downregulated ELC; however, at the gene level this expression profile is arrhythmic (Supporting Figure S5). These findings highlight the importance of performing these analyses at the transcript level.

### Motif enrichment analysis

To identify putative regulators of the pathway-level rhythmicity changes that we observe with age for all ELCs, RLCs and LLCs, we performed a motif enrichment analysis. Briefly, we scanned promoter regions of all transcripts in the transcriptome for known motif instances using the motif scanning tool Find Individual Motif Occurrences (FIMO) [30], and used a hypergeometric test to determine the level of enrichment of each instance among transcripts in DAVID pathway clusters (Figure 7A, workflow diagram). The results of the motif analysis are shown in the heatmap and bar plot in Figure 7A. Plotted clusters are significant (q-value ≤ 0.05) after performing a Benjamini-Hochberg multiple test correction, contain at least ten transcripts, and have a DAVID pathway enrichment score of 1.3 or higher (the full table of motif analysis results is available in Supporting Table S8).

**Figure 7.**
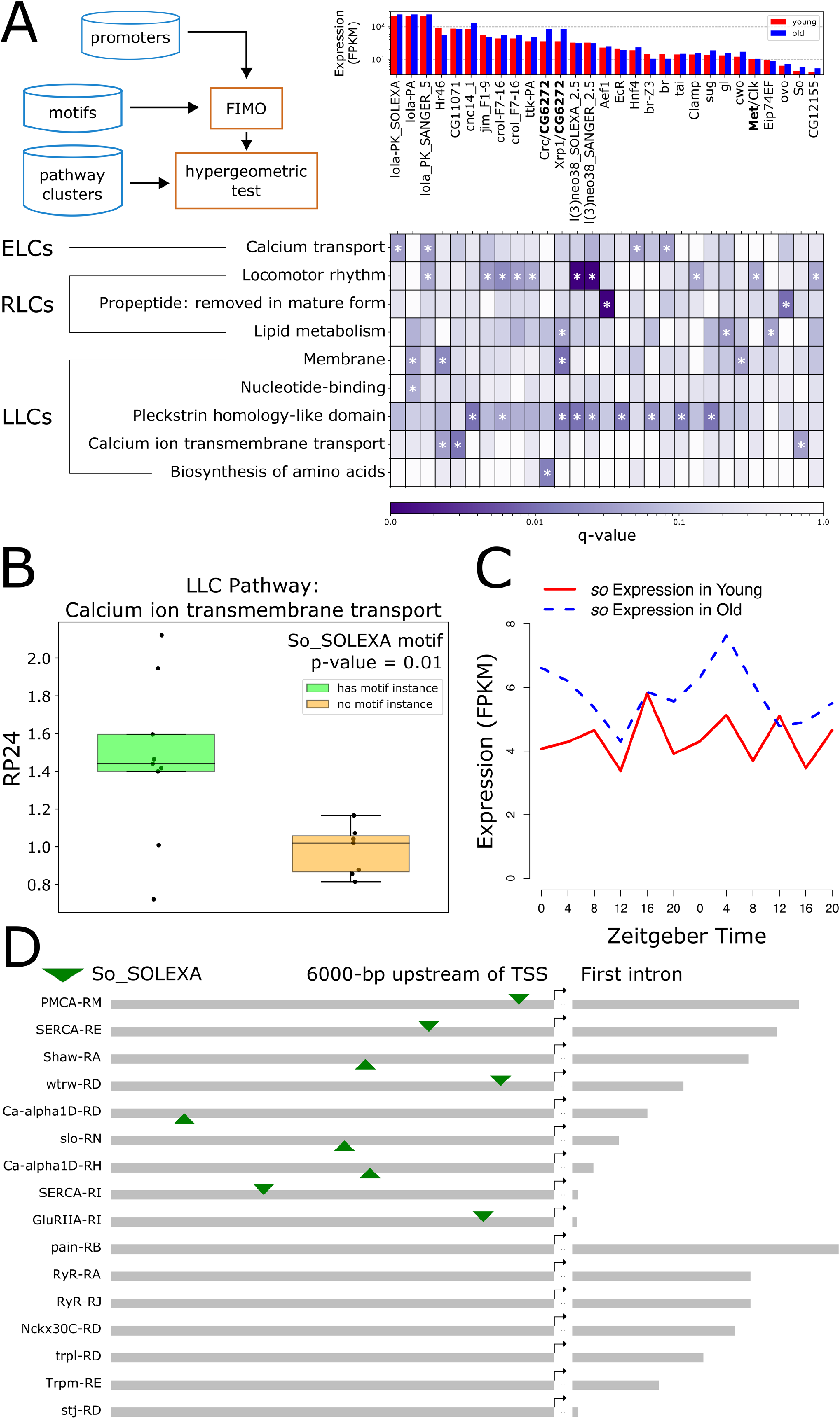
Transcription factor binding site motif analysis of promoter regions. **A**. Heatmap shows enrichment of transcription factor binding site motifs in promoters of transcripts belonging to DAVID clusters. DAVID clusters are on the y-axis, and transcription factors corresponding to binding site motifs are on the x-axis. DAVID clusters shown have at least 10 transcripts, and transcription factors pass an expression threshold of 5 FPKM in young or old flies based on our experimental data. Hue represents q-value on a logarithmic scale. Asterisks mark results with q-value ≤ 0.05. Bar plot shares an x-axis with the heatmap, and shows young (red) and old (blue) expression of each transcription factor. Dimer partners are separated with a forward slash (/); the expression of the dimer partner with the lowest expression (bolded) is represented in the bar plot. Duplicate transcription factor symbols have different motif position weight matrices, and are distinguished by inclusion of sequencing platform information. Diagram of workflow is show in the upper left-hand corner of panel A. **B**. Boxplot compares RP24 (in old flies) for LLC transcripts involved in calcium ion transmembrane transport, with and without significant So transcription factor binding site motifs. **C**. Gene expression profile of *so* in young (red) and old (blue) flies. Lights-on is Zeitgeber Time (ZT) 0, and lights-off is ZT12. **D**. Binding site motif locations for So in promoters of LLC transcripts involved in calcium ion transmembrane transport. Motif locations are denoted by a downward-pointing (forward strand) or upward-pointing (reverse strand) green triangle. Promoters are ordered by length of first intron (grey bar following rightward pointing arrow) within subgroups containing or lacking motif instances for So.

Top results include the enrichment of RLC transcripts belonging to the pathway “propeptide: removed in mature form”, for motif instances of the transcription factor AEF-1. These transcripts all share a non-standard processing pipeline in which a small piece of the protein (the propeptide) is removed during maturation or activation (https://www.uniprot.org/help/propep). RLC transcripts involved in locomotor rhythm were enriched for motif instances for two heterochromatin silencers, l(3)neo38 and CROL [31]. Surprisingly, we found that RLC transcripts involved in locomotor rhythm were also significantly enriched for motif instances for the CLK/Met heterodimer (p-value = 0.0037, q-value = 0.044), rather than the well-known CLK/CYC heterodimer (p-value = 0.096, q-value = 0.26). Both are known to bind to canonical E-box motifs [32], but the flanking bases differ between the two motifs (Supporting Figure S6). Ten of these locomotor rhythm transcripts belong to genes which have previously been shown to be bound by CLK [33], including the core clock genes *per, tim, vri*, and *Pdp1*. We note that clockwork orange (*cwo*), a basic helix-loop-helix transcription factor involved in regulating rhythmic gene expression within the transcriptional feedback loop that keeps circadian time, was also identified as regulating membrane associated proteins.

Transcripts from multiple pathways were enriched in motif instances for Irbp18 heterodimers. Irbp18 is a transcription factor containing a basic leucine zipper domain (BZIP) which is involved in repairing double-stranded DNA breaks induced by transposase enzymes at P-element sites [24] that has been shown to respond to blue light [10]. We found enrichment of heterodimers formed by Irbp18 and two other BZIP transcription factors, CRC and Xrp1. Five of the seven *Xrp1* isoforms are significantly upregulated in our dataset, including the two most highly expressed in old flies (*Xrp1-RD* and *Xrp1-RE*). One of the *Irbp18* isoforms (*Irbp18-RA*) also shows a significant increase in expression level with age (Supporting Figure S7).

LLC transcripts involved in dehydrogenase activity and the biosynthesis of amino acids were enriched for CRC/Irbp18 heterodimer motif instances. Other LLCs, encoding proteins containing Plekstrin homology-like domains and transmembrane domains, were enriched for Xrp1/Irbp18 motifs. RLC transcripts involved in lipid metabolism were also enriched for Xrp1/Irbp18 motifs. In addition, although not significant after multiple test correction (p-value = 0.0057, q-value = 0.14), ELC transcripts in the glutathione metabolism pathway also contained motif instances for the Xrp1/Irbp18 heterodimer.

To explore the effect of putative regulators on expression level and rhythmicity, we tested all significant clusters for differential expression and differential rhythmicity between transcripts with and without significant motif instances using a Welch’s T-test. We found a significant upregulation of rhythmicity in LLC transcripts involved in calcium ion transmembrane transport with the motif instance for the transcription factor So (*sine oculis*) compared to transcripts in this pathway without the motif instance (Figure 7B). Expression data from our experiment shows that the gene encoding this transcription factor, *so*, is also rhythmic in old flies (Figure 7C). We visualized the location of the So motif in the 9 out of 16 (56.25%) transcript promoters in this LLC cluster with the motif instance, and note that several transcripts derive from the same gene product, although they have different promoters (*SERCA-RE* and *SERCA-RI, Ca-alpha1D-RD* and *Ca-alpha1D-RH*) (Figure 7D). We also found a significant reduction of rhythmicity (p-value = 0.042) in RLC transcripts involved in locomotor rhythm with a motif instance for the transcription factor Ttk (*tramtrack*) compared to transcripts in this pathway lacking the motif instance (Supporting Figure S8A). This pathway includes twelve transcripts derived from all the core clock genes; seven of these contain an instance of the Ttk motif (Supporting Figure S8B). Ttk has been linked to the circadian clock as a putative regulator of *pdf*, the main circadian neuropeptide in *Drosophila* [34].

We also performed (as above) motif enrichment analysis on LLC transcripts with phases between ZT2 and ZT6 (full list in Supporting Table S5). The only statistically significant result was for the BZIP heterodimer Xrp1/Irbp18 (Supporting Figure S9). Pathway analysis revealed that LLC transcripts enriched for this motif are involved in alternative splicing, imaginal disc-derived processes (leg morphogenesis and wing vein specification), peripheral nervous system development, and the cell cortex, as well as transcripts encoding proteins containing membrane and IPT (Immunoglobulin-like, Plexins, and Transcription factors) domains. The full list transcripts enriched for Xrp1/Irbp18 motif instances is available in Supporting Table S6, and the full list of pathway analysis results is available in Supporting Table S7.

## Discussion

We performed a systems-level study of age-related diurnal expression in the *Drosophila* transcriptome that revealed important changes in rhythmicity in old flies as well as functionally-related transcripts that remain robustly rhythmic after aging. We developed the RP24 as a useful and interpretable score that better highlights highly-precise rhythms and changes in rhythmicity than previous Fourier-based methods. We show the value in focusing on transcripts as opposed to genes, and note an example that is rhythmic at the transcript level, but not at the gene level.

An important message from our study is that a substantial portion of transcripts rhythmic in young maintain a cycling pattern in old (RLCs) consistent with the persistence of a functional circadian clock. A portion of these transcripts are bound by CLK in young flies [33], including core clock transcripts and *norpA*, which is required for phototransduction [35]. We found RLCs to be enriched for light- and circadian-associated transcripts, as well as for transcripts encoding mitochondrial ribosomal proteins. Previous studies have shown that mitochondrial proteins accumulate rhythmically and at the same time, including several metabolic genes, but comparisons of RNA and protein expression did not show correlation [36]. The strong in-phase rhythmicity of transcriptions involved in mitochondrial translation throughout lifespan suggests a mechanism for coordination of these mitochondrial proteins. Although many transcripts related to mitochondrial function remain rhythmic after aging, others lose rhythmicity in old flies (Figure 4). We show that a similar pattern occurs with transcripts involved in phototransduction (Figure 5). The observed loss of pathway coordination after aging suggests that subsets of transcripts within a pathway (e.g. mitochondrial function) have different or additional regulatory inputs that determine whether they remain rhythmic after aging. Changes in expression of regulatory factors after aging may contribute to transcript expression changes that result in pathway dysregulation and consequently some observed detrimental age-related loss of rhythmicity. We show that the core clock transcripts remain rhythmic, but have reduced RP24 scores in old flies when binding site motifs for the transcription factor encoded by the age-upregulated *ttk* gene are present in their promoters. This is perhaps one example of the age-onset “rewiring” of the diurnal regulatory network.

Our study detected age-related changes in phase of oscillatory genes. While transcripts belonging to the core circadian clock machinery remain rhythmic in old flies, they undergo phase advances with age. Studies in human subjects have also observed phase advances in older individuals, while the circadian period remains unchanged [37]. In contrast to clock genes, we show a global transcriptomic trend toward a phase delay. Transcripts rhythmic only in young flies (ELCs and RLCs in young) predominantly peak before lights on, while transcripts rhythmic only in old flies (LLCs and RLCs in old) tend to peak after lights on (ZT2 to ZT6 for LLCs). The predominant phase of LLCs being 2-6 hours after lights on suggests the possibility of a light-activated mechanism.

We show that the promoter regions of LLCs with a phase from ZT2 to ZT6 are significantly enriched for motif instances of the transcription factor heterodimer Xrp1/Irbp18. Xrp1 and Irbp18 are BZIP transcription factors that heterodimerize to repair DNA double-stranded breaks caused by transposases near P-element sites [24], and transcripts for both genes are upregulated in old flies in our dataset. Both of these genes have previously been shown to be upregulated in response to blue light [10], which induces expression of stress-response genes [9-11]. We suggest a program by which flies respond to accumulated light-induced stress with increased expression of DNA damage repair regulators Xrp1 and Irbp18, which regulate the expression of LLCs peaking from ZT2 to ZT6. Irbp18 is orthologous to CCAAT-enhancer binding protein gamma [24], one of the human CCAAT-Enhancer Binding Proteins (C/EBP). Xrp1 also has high sequence similarity to mammalian C/EBP transcription factors [38], making the study of the putative regulation of light-activated transcripts by this heterodimer an important topic for future study.

We detected a prominent shift in glutathione-related metabolism with age, including a loss of rhythmicity and an upregulation in transcripts encoding several glutathione-S-transferases, and a gain of rhythmicity in transcripts encoding the modifier subunit (*Gclm-RA* and *Gclm-RB*) and catalytic subunit (*Gclc-RB*) of glutamate cysteine ligase (GCL), the rate-limiting enzyme in glutathione biosynthesis. This is consistent with our previous studies reporting circadian oscillations in *Gclc* gene expression in young flies and associated rhythmic GCL protein level and enzymatic activity [39]. However, in old flies these rhythms were abolished and both *Gclc* expression and GCL activity were at constantly high levels [40]. This suggests that in addition to CLK/CYC, other transcription factors contribute to the regulation of these genes in old flies. Our motif analysis revealed an enrichment for Xrp1/Irbp18 heterodimer binding site motifs in promoters of glutathione metabolism transcripts, suggesting a potential alternate regulator of this pathway during aging. Xrp1 has recently been shown to induce glutathione-S-transferases as part of the PERK-mediated unfolded protein response [41], corroborating our motif analysis results.

Our results showing gain of transcript rhythmicity with aging are consistent with other studies [7, 8, 12]. We analyzed promoters of LLC transcripts grouped by function for transcription factor binding site motifs and identified sine oculis as a putative regulator of the 16 transcripts annotated as involved in calcium ion transmembrane transport. The gene encoding this transcription factor (*so*) is rhythmic in old flies in our dataset, consistent with a recent single-cell transcriptomics study which found *so* to be rhythmic in a subset of *Drosophila* circadian clock neurons [42]. This suggests that sine oculis may be a regulator of age-dependent rhythmicity in these LLC transcripts.

We found a cluster of 32 LLC transcripts with annotations related to metabolism including the KEGG pathway “Biosynthesis of amino acids”, and several genes encoding dehydrogenase enzymes (*Gdh, Ldh, Aldh, Adh, SdhA*). It is tempting to speculate that increased rhythmic expression is protective. However, functional analysis of one of the most robustly rhythmic LLCs, lactate dehydrogenase (*Ldh*, formerly *ImpL3*), determined that its high gene expression accelerates the aging process, while reduced expression delays aging [26]. *Ldh* expression increases in old flies during the light phase but remains low in constant darkness, which also extends fly lifespan [11]. Taken together, these results show that conclusions regarding protective or detrimental effects of elevated expression of a given gene require experimental verification.

Finally, this study demonstrates the value of flies subjected to light/dark cycles as a model for human exposure to environmental light sources. Humans are exposed to increased blue light, both in duration and intensity, from ever-present artificial light emanating from LED and other screens. Flies have been used to examine the effects of blue light on longevity [11], with gene-expression-level results applicable to human health and disease. This study uses flies as a model to understand diurnal changes at the transcriptome level during natural aging, and increases our knowledge base for addressing age-related concerns in the aging human population.

## Methods

### Read alignment and quantification

Raw RNA-seq reads from head samples of *white* flies collected at 4h intervals around the clock were sequenced and preprocessed as previously described [8]. Data are accessible at the GEO accession GSE81100. Quality filtered and trimmed reads were aligned to the *Drosophila melanogaster* genome (BDGP release 6.21/dm6) using hisat2 version 2.1.0 [43] using the parameters “--max-intronlen 10000 --rna-strandness F”. Cuffdiff [44] was used to quantify transcript abundance in Fragments per Kilobase per Million mapped reads (FPKM). Only protein-coding transcripts were included in downstream analyses.

### Detection of transcript rhythmicity with RP24

Our analysis of transcript expression profiles considers expression as a function of time as *E*(*t*) sampled discretely at *N* time points in increments of Δ*t* such that *t*_*n*_ = *n*Δ*t* over a span of *T* hours. In our data, *T* = 48 hours, and we sampled every 4 hours for a total of *N*=12 time points. We can then define a discrete series of expression values at these times as *E*_*n*_ = *E*(*t*_*n*_). The Fourier transform for the expression time series is computed from

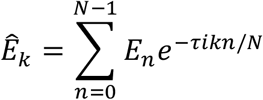

where *τ* = 2*π*. The Fourier transform decomposes the expression profile into sinusoidal waves with a period *T*_*k*_ = *T*/*k*. It is useful to consider the power spectrum *P*(*T*_*k*_) that defines the strength of contribution of a particular period *T*_*k*_.

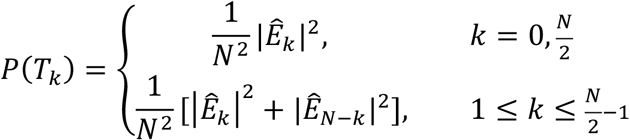

To detect the period of 24 hours (*T*_2_ = 24) that are relevant for circadian rhythms, we define RP24 as the relative power of the 24-hour period compared to other periods to quantify rhythmicity:

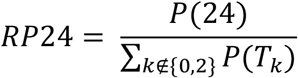

### Calculation of other RPT_k_ values

The RP24 is generalized in Supporting Figure S1 to other Fourier periods 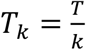 using the discrete Fourier power spectrum. These non-24-hour periods can be measured analogously to the RP24 score where a different period is deemed the signal rather than the noise.

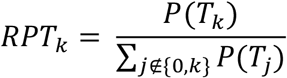

This includes periods corresponding to frequencies up to and including the Nyquist frequency, or half the sampling rate. In our case, with 12 time points over 48 hours, the Nyquist frequency is 6. Thus, dividing 48 hours by frequencies of 1-6 yields the valid periods for which an RP score can be measured: 48, 24, 16, 12, 9.6, and 8.

### Simulated Expression Profiles

We simulated some expression profiles for Supporting Figure S2. These expression profiles were based on the sinusoidal equation,

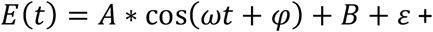

where *E*(*t*) is expression of simulated transcript, and *t* is time of day. The time values were set at intervals every 4 hours between 0 and 24 (0, 4, 8, 12, 16, 20, 24) similar to the RNA expression data we collected. Amplitude was held constant at 1, with omega fixed at 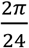 corresponding to 24-hour periods, *φ* uniformly ranged from 0 to 2*π*, and *B* ranged from 0 to 300. To generate the simulated transcript expression data, a level of noise *ε* ∼*N*(0, *σ*^2^) was added to the expression. Simulated transcripts were set with a signal to noise ratio (SNR) ranging between 0.01 and 15. The SNR value of each transcript was applied to set the level of noise (*σ*), through the following equation 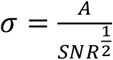. The level of noise added to each calculated expression value was randomly sampled from a normal distribution.

### Analysis of phase changes

While we describe the expression of the 24-hour component as *E*_24_(*t*) = *A* cos((2*π*/24*hr*)*t* + *φ*), the phase is defined as the time of peak expression corresponding to the input to the cosine function to be 0, so that:

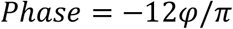

This quantity can be calculated for each transcript from the 24-hour Fourier coefficient regardless of rhythmicity category, but is confidently accurate for transcripts with significant RP24 values.

### Dot and Arrow Plots

The dot and arrow plots describe the change in the state of rhythmicity collection of genes or transcripts. The state of rhythmicity refers to the combination of *RP*24 and the *Phase*. The distance from the origin is computed as *RP*24 with a sigmoidal scaling that maps to a number between zero and one:

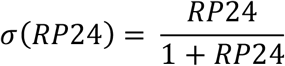

For the *S* = *log*_2_(*RP*24), this transformation is represented as a base-2 logistic function

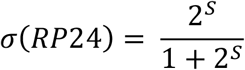

The phase is shown as the angle relative to the positive y-axis, corresponding to ZT0 on a 24-hour clock. To achieve this, we plot a complex number in polar coordinates, and *Phase* is mapped back to radians, and rotated 90° or 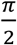 radians by adding 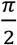. This corresponds to the fact that normally an angle of 0° corresponds to the positive x-axis.

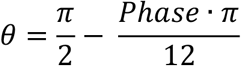

Both the dot and the location of the arrow are then defined as a point on the complex plane computed from

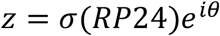

using the *RP*24 and *θ* from young for the dot, and from old for the location to which the arrow points.

### Pathway analysis

We performed pathway analysis on ELCs, RLCs, and LLCs using the DAVID Functional Annotation Tool version 6.8 [22]. DAVID produces clusters of transcripts with an enrichment score S, defined as the −*log*_10_() of the geometric mean of the q-values for each cluster annotation. We required an enrichment score threshold of at least 1 and at least 10 transcripts per cluster for inclusion in the plot in Figure 4A.

### Differential expression analysis

We used Cuffdiff [44] to test for differential expression of transcripts between young and old flies independent of time of day. All samples corresponding to young flies (all replicates, all time points) were used as the first condition, and all samples corresponding to old flies were used as the second condition. We considered transcripts significantly differentially expressed which passed a q-value threshold of 0.05.

### Motif analysis

We analyzed data from the *Drosophila* transcriptional regulatory element database (RedFly) to determine an appropriate window size for the promoter region upstream of the transcription start site for each transcript [45] (Supporting Figure S10). Based on our analysis and previous reports in the literature, we defined promoter regions as the 6000-bp region upstream of the annotated transcription start site for each transcript, plus the first intron. These two regions were concatenated with a 20 base pair linker of Ns to prevent false identification of motifs spanning the junction between regions.

We compiled a list of transcription factor binding site motifs from two publicly available databases: FlyFactorSurvey [46] and JASPAR [47]. In addition, we used MEME-ChIP [48] for *de novo* identification of cnc binding motifs from four ChIP-seq results files [49], using the parameters “-meme-mod zoops” for each file. The six significant (E-value < 1) motifs we identified were included in the motif analysis. To filter out low-quality motifs and to restrict our analysis to transcription factors expressed in our dataset, we required all motif instances to pass p-value threshold of 0.00005, and the corresponding transcription factors a median FPKM threshold of 1. We tested for enrichment in the number of promoters with occurrences of the resulting 629 known transcription factor binding motifs in the promoters of transcript groups of interest. FIMO [30] was used to scan for motifs in the promoter regions of all transcripts in the transcriptome using the parameters “--no-qvalue --thresh 1e-4”.

We applied a more stringent DAVID cluster enrichment score of −*log*_10_(0.05) ≈ 1.3 to correspond to a geometric mean of multiple-test corrected p-values of 0.05. We also required at least ten transcripts per cluster. We used a hypergeometric test to determine the enrichment of the motifs in promoter regions of transcripts belonging to each these DAVID clusters compared to the rate of motif occurrence in the global set of promoters for all transcripts in the transcriptome. We applied a Benjamini-Hochberg FDR multiple test correction to the resulting p-values for each cluster.

### Filtering out transcripts with expression “spikes”

We detected several transcripts that exhibited sharp spikes of expression at regular intervals. Although these spikes occur at regular intervals, they were not detected by other Fourier-based approaches. It is well known that “spiky” expression patterns are not easily detected by these methods [15]. Because we were unable to validate these transcript expression profiles with qPCR (data not shown), we filtered them out of our analysis.

To detect expression profiles that spike up periodically at one measured time point, we define a baseline expression level *E*_0_, plus a burst of expression Δ*E* at a particular time point *φ*, which is analogous to the phase:

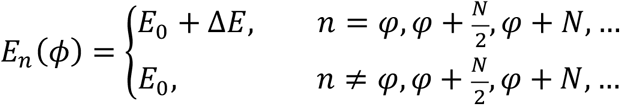

The Fourier transform for these transcripts does not show a large RP24 or low p-value from our methods. Because we could not validate their expression, we developed methods for detecting these transcripts, and removed transcripts with these rhythms from our groups of rhythmicity changes. We refer to these rhythmic bursts of expression as “staccato”.

The Fourier transform for these transcripts can be computed as follows:

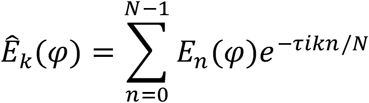

Expanding this sum gives

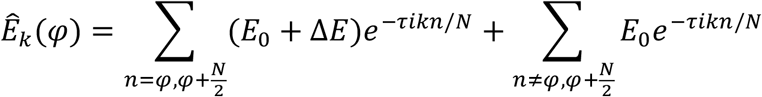

Grouping terms gives the expression

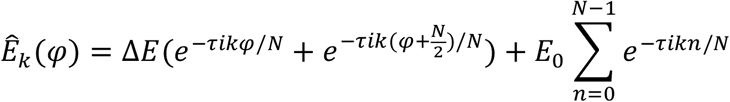

Which can be reduced to

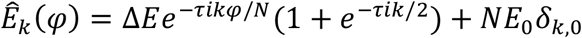

Therefore, the Fourier coefficients reduce to the following three cases:

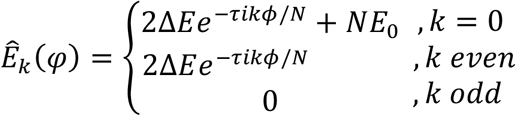

Therefore, idealized staccato rhythms should have non-zero, even-numbered Fourier coefficients. Real data can be variable, which led us to define a score called the “spectral parity,” quantifying the extent to which the Fourier coefficients are biased toward even-numbered values.

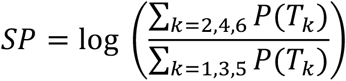

We also observed that the argument of the even-numbered Fourier coefficients, *arg*(*Ê*_*k*_ (*ϕ*)), which corresponds to the phase of the sinusoidal wave, should be equal. This led us to define a score called the “phase variance of even coefficients,” defining the degree of dispersion of the phases of the even-numbered Fourier coefficients.

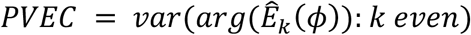

When the PVEC is low, the even terms are coordinated and reinforce the spiked expression pattern.

## Author Contributions

B.S. and D.A.H. conceived of the method, B.S., R.M.F., P.M., B.L., and D.A.H. performed the formal analyses, and B.S., R.M.F., J.M.G., and D.A.H. wrote the paper.

## Acknowledgements

Barbara Gvakharia for help with investigation of mitochondrial ribosomal protein literature. Eileen S. Chow for conducting qPCR experiments.

## Code Availability

Code for this project is available at https://github.com/hendrixlab/RP24.

