## Supplementary Figures for "Discovery and Visualization of Age-dependent Patterns in the Diurnal Transcriptome of Drosophila"

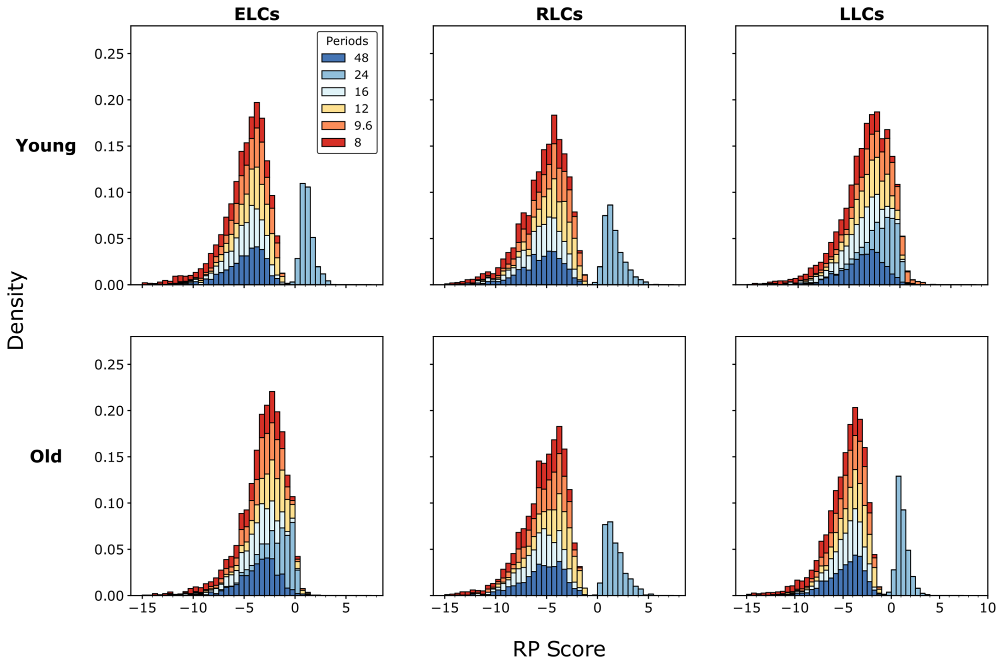


Supporting Figure S1: Distributions for the relative power for different Fourier periods $T_k$ for $k = 1,2,3,4,5,6$.

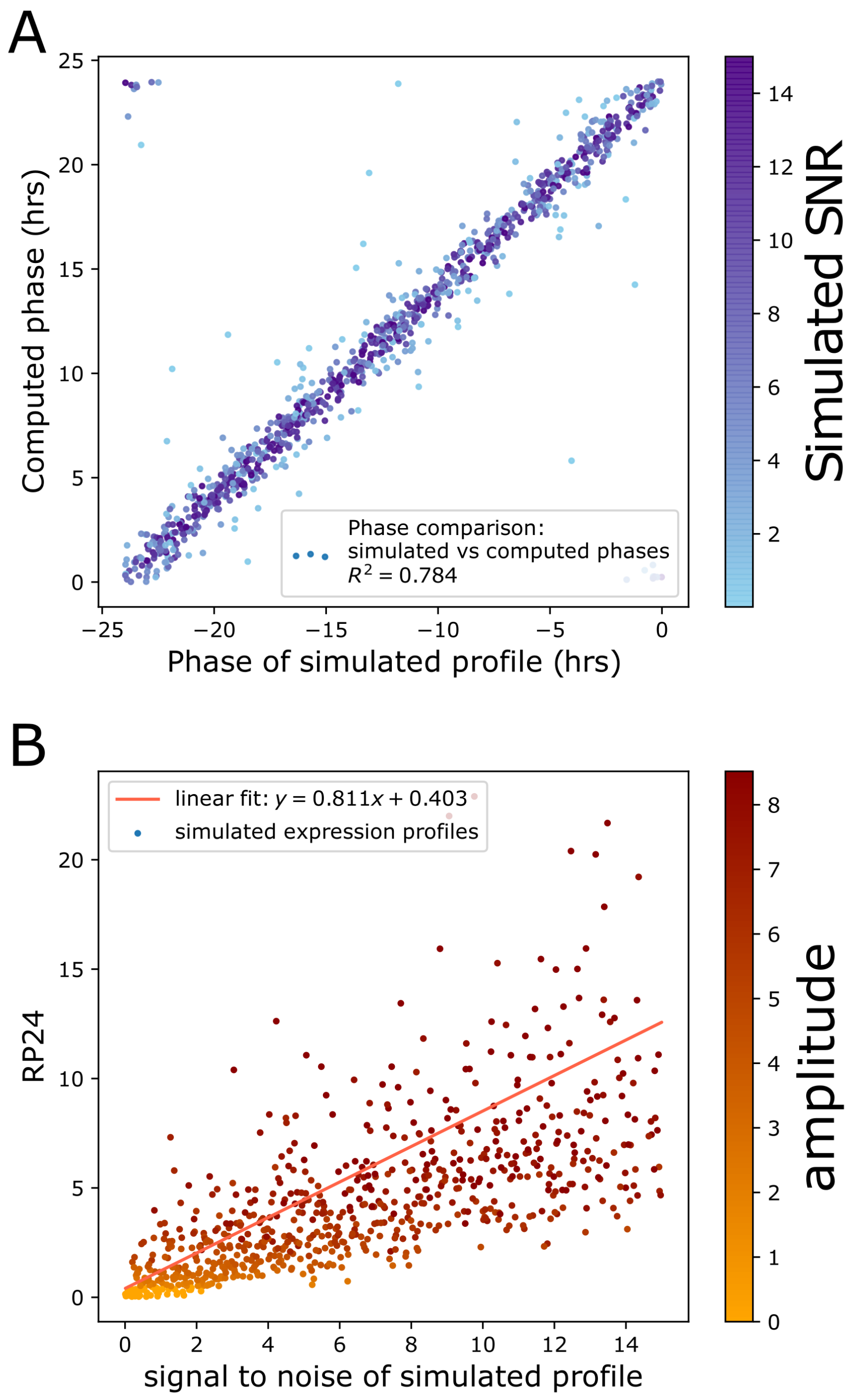


Supporting Figure S2: Phase simulations. **A.** Computed phases for simulated expression profiles show a similar trend as the phase used in the simulation. The values in the upper left are still valid because ZT24 is the same as ZT0. Colors correspond to the signal to noise ratio (SNR) of the simulated expression profile. **B.** RP24 values computed from simulated expression profiles are correlated with the SNR of the simulated expression. Colors correspond to the amplitude of the expression profile.

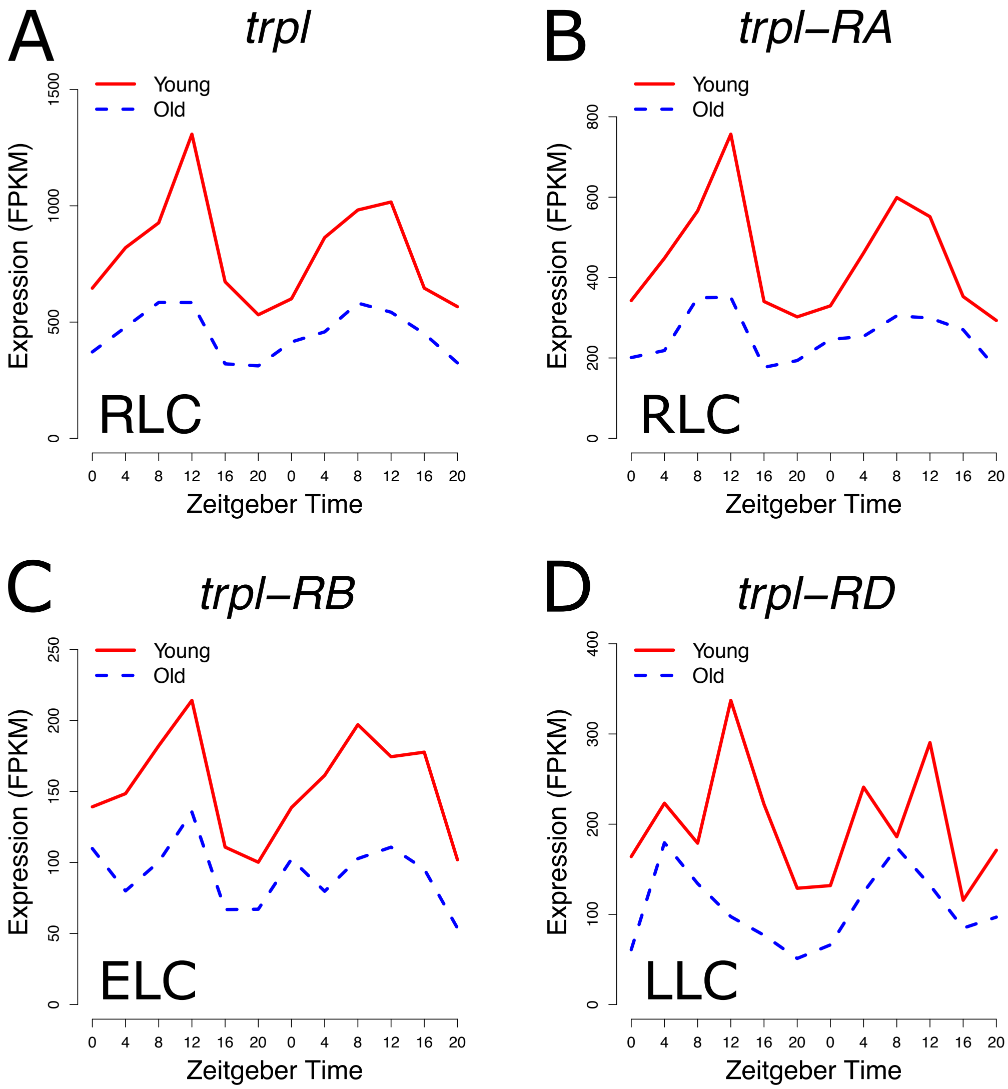


Supporting Figure S3: RNA-seq expression profiles for eye-enriched *trpl*, which is required for phototransduction. Each Zeitgeber time point on the x-axis represents a separate biological replicate. Transcript Rhythmicity group classification is denoted in the lower left-hand corner of each plot. **A.** Gene *trpl* and **B.** transcript *trpl-RA* are rhythmic in both young and old flies. **C.** Transcript *trpl-RB* is rhythmic in young flies only. **D.** Transcript *trpl-RD* is rhythmic in old flies only.

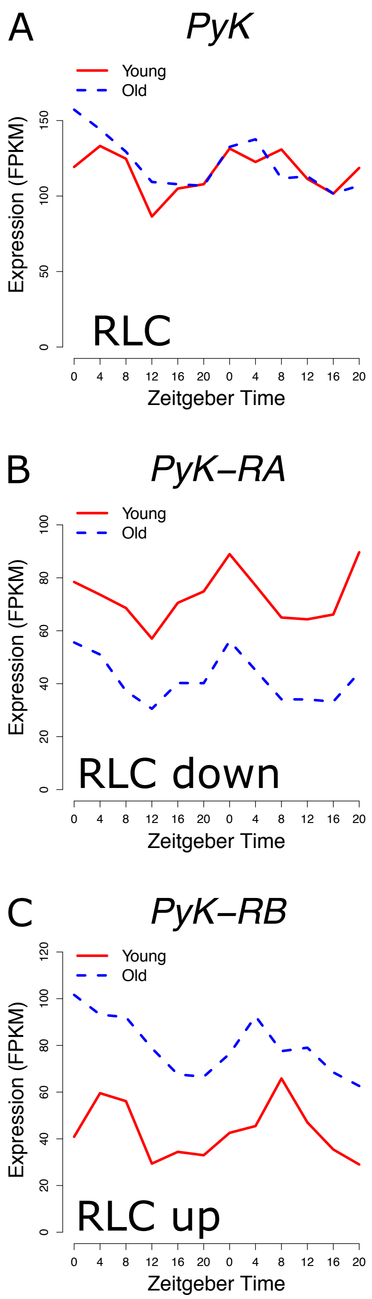


Supporting Figure S4: RNA-seq expression profiles for *PyK*, which encodes pyruvate kinase. Each Zeitgeber time point on the x-axis represents a separate biological replicate. Rhythmicity group classification is denoted in the lower left-hand corner of each plot. **A.** The gene *PyK* is an RLC which does not significantly change expression level after aging. **B.** The transcript *PyK-RA* is an RLC which is significantly downregulated in old flies. **C.** The transcript *PyK-RB* is an RLC which is significantly upregulated in old flies.

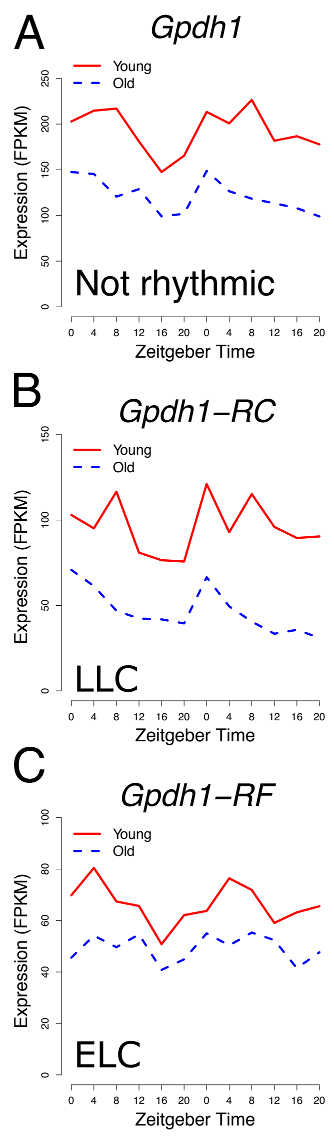


Supporting Figure S5: RNA-seq expression profiles of *Gpdh1*, which encodes for glycerol-3-phosphate dehydrogenase. Each Zeitgeber time point on the x-axis represents a separate biological replicate. Rhythmicity group classification is denoted in the lower left-hand corner of each plot. **A.** The gene *Gpdh1* is not rhythmic in young or old flies. **B.** The transcript *Gpdh1-RC* is rhythmic in old flies. **C.** The transcript *Gpdh1-RF* is rhythmic in young flies.

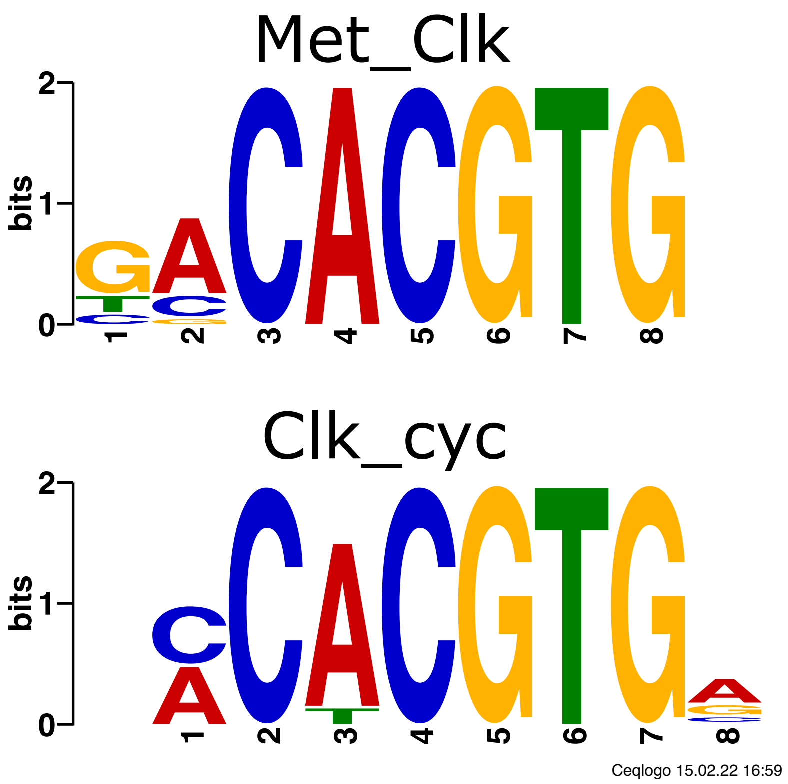


Supporting Figure S6: Logos for motifs for Met/Clk and Clk/CYC heterodimers, created with Ceqlogo.

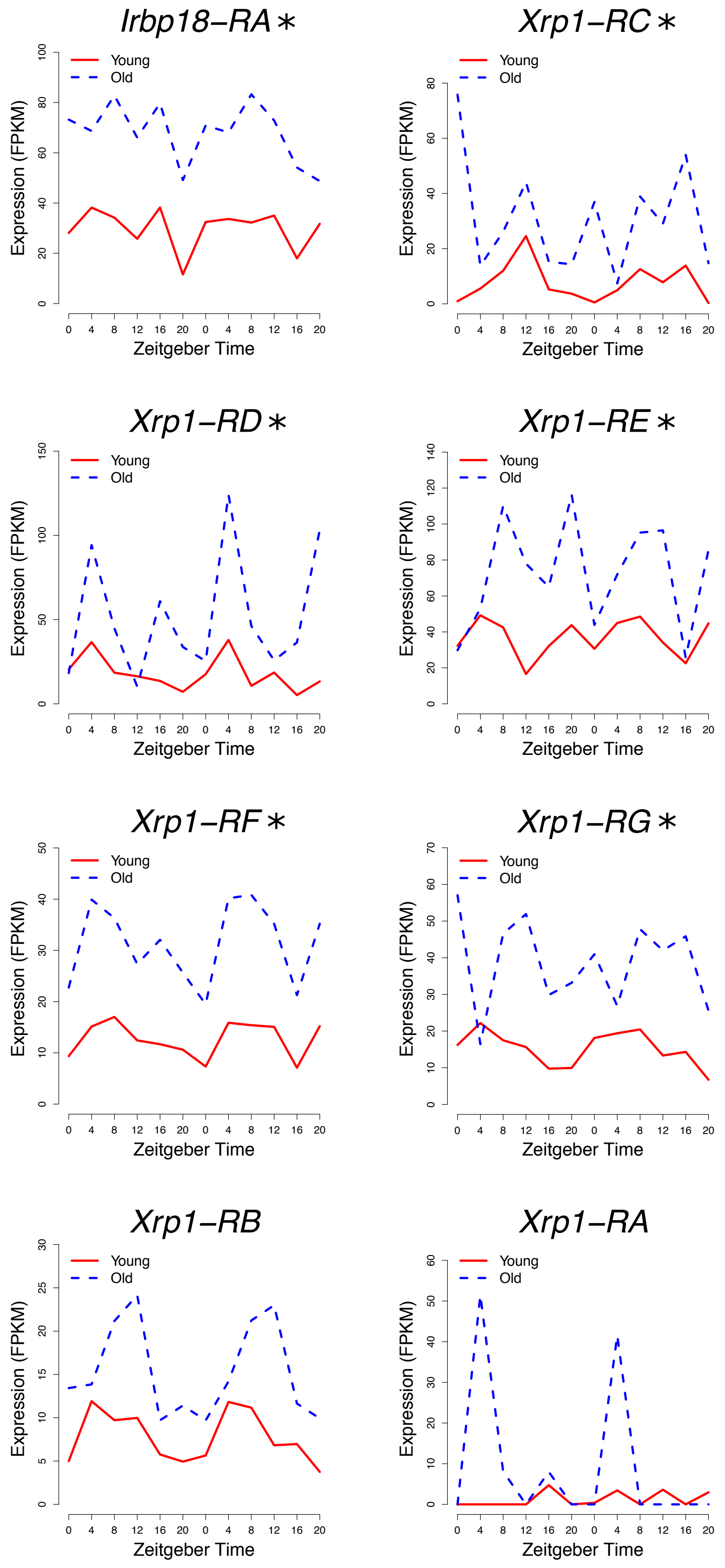


Supporting Figure S7: RNA-seq expression profiles for *Irbp18* and *Xrp1*. Each Zeitgeber time point on the x-axis represents a separate biological replicate. Isoforms marked with an asterisk are significantly upregulated in old flies compared to young.

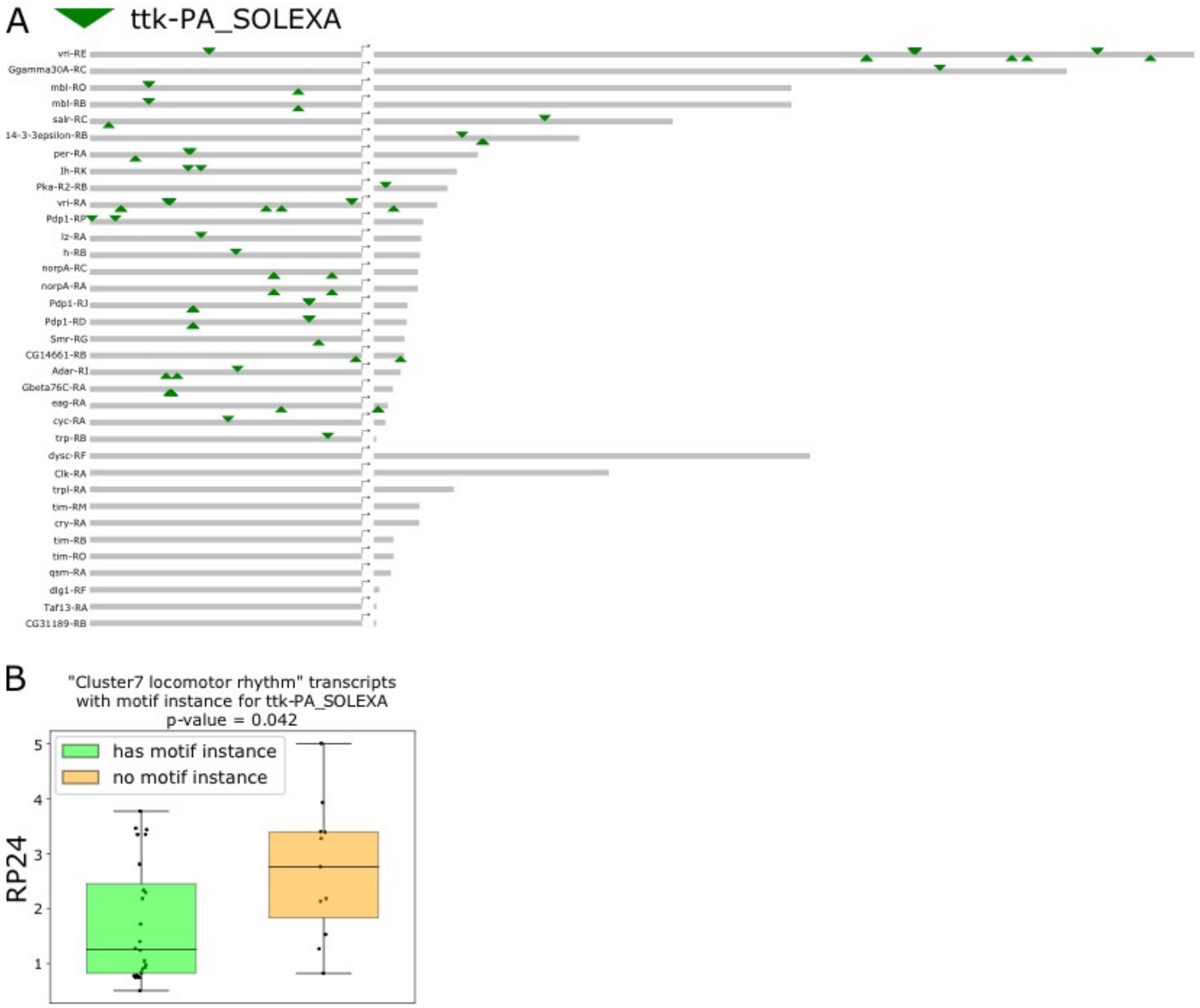


Supporting Figure S8: Locomotor rhythm RLC transcript promotors are enriched for motif instances of ttk binding sites. **A.** Representation of promotor regions and first introns for RLC transcripts in the locomotor rhythm pathway. Isoform name on the left is followed by the 6000 base pair promoter region upstream of the transcription start site, represented by a right-pointing arrow, followed by the first intron of the transcript (variable lengths). Binding sites are denoted with a green triangle; triangles pointing down indicate motif instances on the forward strand, while triangles pointing up indicate motif instances on the reverse strand. Isoforms are arranged in order of decreasing intron length for isoforms with and without ttk binding site motifs. **B.** Comparison of RP24 scores of transcripts with (green) and without (orange) ttk motif instances.

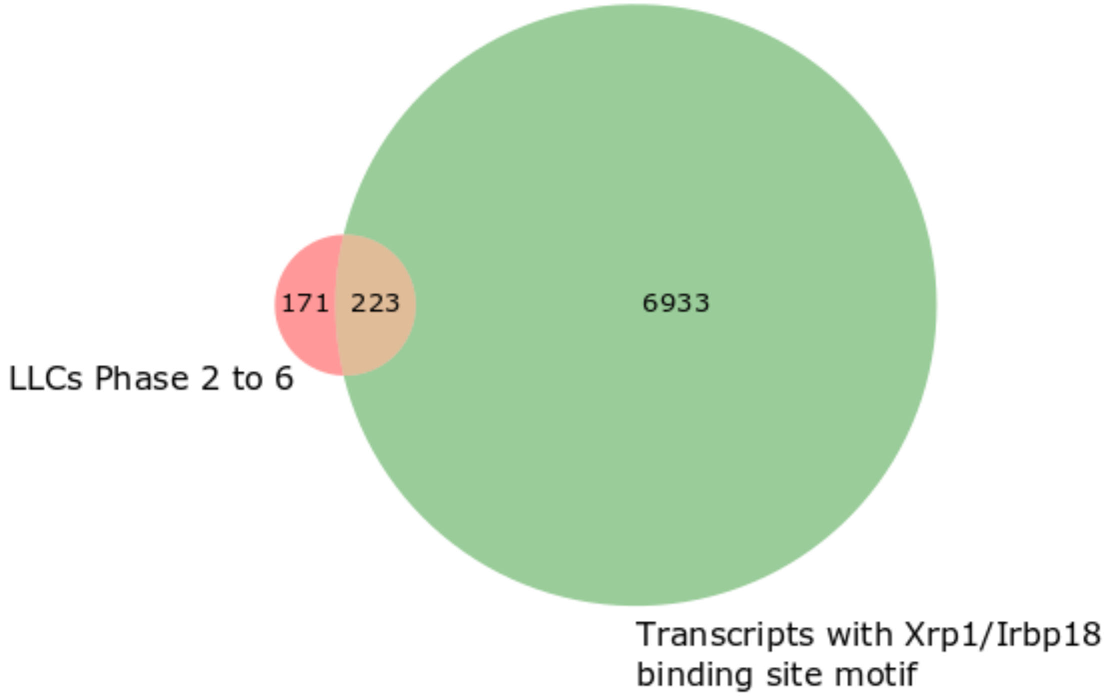


Supporting Figure S9: Venn diagram shows the overlap between LLCs with a phase between ZT2 and ZT6 (red) and all transcripts with a binding site for Xrp1/Irbp18 in the promoter region.

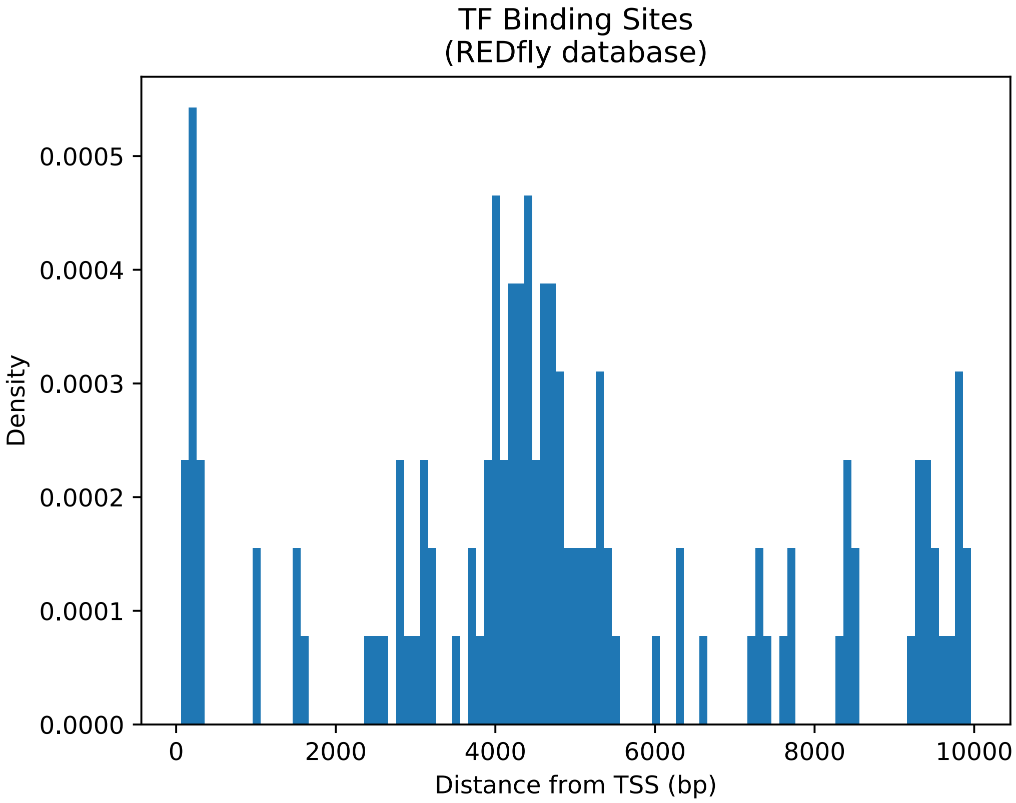


Supporting Figure S10: Density of transcription factor binding sites plotted against distance in base pairs (bp) from the transcription start site (TSS) of nearest gene. Data is from the Regulatory Element Database for Drosophila (REDfly).
